# Characterization of novel and essential kinetoplast-associated proteins in *Trypanosoma brucei*

**DOI:** 10.1101/2024.04.22.590512

**Authors:** Lawrence Rudy Cadena, Michaela Svobodová, Corinna Benz, Vendula Rašková, Ľubomíra Chmelová, Ingrid Škodová-Sveráková, Vyacheslav Yurchenko, Julius Lukeš, Michael Hammond, Ignacio Miguel Durante

## Abstract

The kinetoplast is one of the defining features of kinetoplastid protists and represents a unique concentration of mitochondrial DNA. This subcellular structure is a highly complex assembly of thousands of mutually catenated, circular DNA molecules as well as up to one hundred dedicated proteins. These components work in tandem to replicate and segregate the mitochondrial genome during cellular division, additionally coordinating with the basal body and flagellum through the tripartite attachment complex (TAC) superstructure. Here, we screened the MitoTag localization repository and identified a number of previously undescribed hypothetical proteins exhibiting putative signals within the kinetoplast of *Trypanosoma brucei*. Through endogenous tagging we verify their association with the kinetoplast or TAC. The essentiality for several of these kinetoplast proteins (KP) was assessed by RNAi knock-downs, revealing that the newly characterized KP56, KP84 and KP86 are indispensable for growth of the procyclic stage. Additionally, KP37, KP56, and KP84 displayed alterations in the abundance of maxicircles or minicircles, while the depletion of KP84 and KP86 resulted in cell cycle alternations. Pulldown assays using the endogenously V5-tagged cell lines identified novel interactors, which were additionally subjected to endogenous tagging for subcellular localization, revealing two additional proteins (KP45 and KP66) with dual localization to the kinetoplast and throughout the mitochondrial lumen. This work represents the most extensive identification of novel KPs to date and provides a methodological pipeline for the characterization of remaining KPs to further understand this intricate molecular structure.

## INTRODUCTION

The kinetoplast, one of the defining features of the predominantly parasitic protistan order Kinetoplastea (Alexei Y. Kostygov et al., 2021), represents a complex organelle substructure containing the mitochondrial genome, termed kinetoplast (k) DNA (Jensen & Englund, 2012). The kDNA is a single catenane composed of circular DNA molecules of two types, referred to as maxicircles and minicircles, located within this defined kinetoplast region of the mitochondrial matrix, proximal to the basal body of the flagellum (Maslov, 2019). In the best studied representative, *Trypanosoma brucei*, about two dozen maxicircles (∼23 kb long) and approximately 5,000 minicircles (∼1.0 kb long) are organized into a densely packed kDNA disc (Povelones, 2014). The kDNA maxicircles are homologs of standard mitochondrial genomes and, in *T. brucei*, encode 18 structural genes comprising subunits of respiratory complexes, one mitochondrial ribosomal protein and two mito-ribosomal RNAs (Alfonzo, Thiemann, & Simpson, 1997). The complexity of the mitochondrial genome and transcriptome in kinetoplastid flagellates is further enhanced by the uridine insertion/deletion type of RNA editing, mediated by minicircle-encoded guide RNAs, upon numerous maxicircle-encoded transcripts prior to their translation (K. D. Stuart, Schnaufer, Ernst, & Panigrahi, 2005).

The kDNA disc is secured to the anterior end of the mitochondrion by the tripartite <u>a</u>ttachment <u>c</u>omplex (TAC), a series of filaments which extend through the outer and inner mitochondrial membranes and attach to the basal body of the single flagellum. The TAC is additionally involved in the coordinated replication of the kinetoplast and flagellum (Aeschlimann, Stettler, & Schneider, 2023). During cell division, TAC filaments physically link the replicated kDNA discs to the basal and pro-basal bodies across both mitochondrial membranes, thus tightly coupling their segregation with that of the flagellum (Hoffmann et al., 2018).

The faithful replication of each circle into the daughter kDNA discs occurs via the decatenation of minicircles from the network by the action of disc topoisomerases and their replication in the <u>k</u>ineto-flagellar <u>z</u>one (KFZ), (Jensen & Englund, 2012; Shlomai, 2004), while a recently proposed model posits a separate replication of minicircle subsets at the two <u>a</u>nti<u>p</u>odal <u>s</u>ites (APS), which are large conglomerates of kDNA-associated proteins (Amodeo, Bregy, & Ochsenreiter, 2022). Mechanisms governing maxicircles replication are less clear, but DNA rings have been observed remaining in the center of the kDNA disc (Carpenter & Englund, 1995), forming a transient thread during segregation called the ‘nabelschnur’ (Jensen & Englund, 2012).

At least 57 proteins, many of which are confined to and conserved across kinetoplastid flagellates, are currently known to comprise the kinetoplast of model species *T. brucei*. These proteins typically play roles in maintaining structural integrity of the kDNA, as well as its replication and segregation, which precedes cell division (Amodeo et al., 2022; Jensen & Englund, 2012; Schneider & Ochsenreiter, 2018). Among these proteins, the gradual release of minicircles by topoisomerase II activity is essential for their replication (Englund, 1979), while helicase TbPIF2 is required for maxicircle replication (Lindsay, Gluenz, Gull, & Englund, 2008; Liu et al., 2009).

The correct implementation of replication-associated processes for the sole repository of genetic information within the singular mitochondrion of *T. brucei* is essential for cell viability. It remains to be established how the elaborate kDNA network can reliably partition, particularly since some essential minicircles are present in just a few copies amongst the ∼10,000 duplicated minicircles prior to division (Gerasimov et al., 2021; S. J. Li et al., 2020; Meehan et al., 2023).

The complexity of the kDNA network invites speculation as to how and why such a critical structure could have arisen within the common ancestor of kinetoplastid flagellates (Lukes et al., 2002). Equally, proteins constituting the kinetoplast have long been regarded as attractive targets for drug development, since kinetoplastid parasites are causative agents of a range of serious diseases of mammals including humans (K. Stuart et al., 2008).

A majority of the 8,721 proteins encoded by the *T. brucei* genome were recently tagged and sub-localized to specific cell compartments in frame of the TrypTag project (Billington et al., 2023). The investigation of mitochondrial proteins from this study, referred to as MitoTag, identified 1,239 proteins localized into the single reticulated mitochondrion, including 337 previously unidentified proteins, some of which co-localized with the kinetoplast (Pyrih et al., 2023), representing candidates of interest. Accordingly, we initiated a follow-up investigation to validate the status of these putative kinetoplast-associated proteins.

Here, we describe *via* several approaches a total of eight proteins as genuine components of the kinetoplast, with two of these proteins appearing to constitute novel TAC components. Moreover, we demonstrate that several of these proteins are essential for cell viability and that their ablation results in both characteristic and novel phenotypes associated with kDNA.

## RESULTS

### Validation of novel kinetoplast-associated proteins

Nine kinetoplast candidate proteins were selected for analysis in the procyclic stage of *T. brucei*, four of which (Tb927.6.4510, Tb927.7.4810, Tb927.7.5320 and Tb927.9.13820) were newly annotated as localizing to this structure in MitoTag (Pyrih et al., 2023). Five other proteins (Tb927.4.2780, Tb927.8.4040, Tb927.10.15660, Tb927.11.2360 and Tb927.11.14120) were annotated in MitoTag as exhibiting an artificial <u>k</u>inetoplast <u>p</u>roximal <u>e</u>nrichment (KPE) localization, but upon our closer inspection, appeared to be genuine components of the kinetoplast.

In order to establish whether these candidate proteins constitute genuine components of the kinetoplast, rather than mitochondrial matrix proteins exhibiting the KPE as a consequence of endogenous tagging by mNeonGreen (Pyrih et al., 2023), we generated cell lines in which individual genes were tagged via the V5 epitope. The resulting endogenously tagged cells were subjected to subcellular fractionation to first confirm that the targeted proteins co-fractionated with employed mitochondrial markers. Immunofluorescence (IFA) analysis was then employed to determine if the proteins in question co-localized with the kinetoplast signal, or if another localization signal was observed. Proteins that localized to the kinetoplast were subjected to further investigation **(Fig. 1)**, whereby their essentiality was addressed by RNAi knock-down studies, and their endogenous V5 tags were used as bait to identify co-precipitating proteins and thus new kinetoplast candidates.

**Figure 1.**
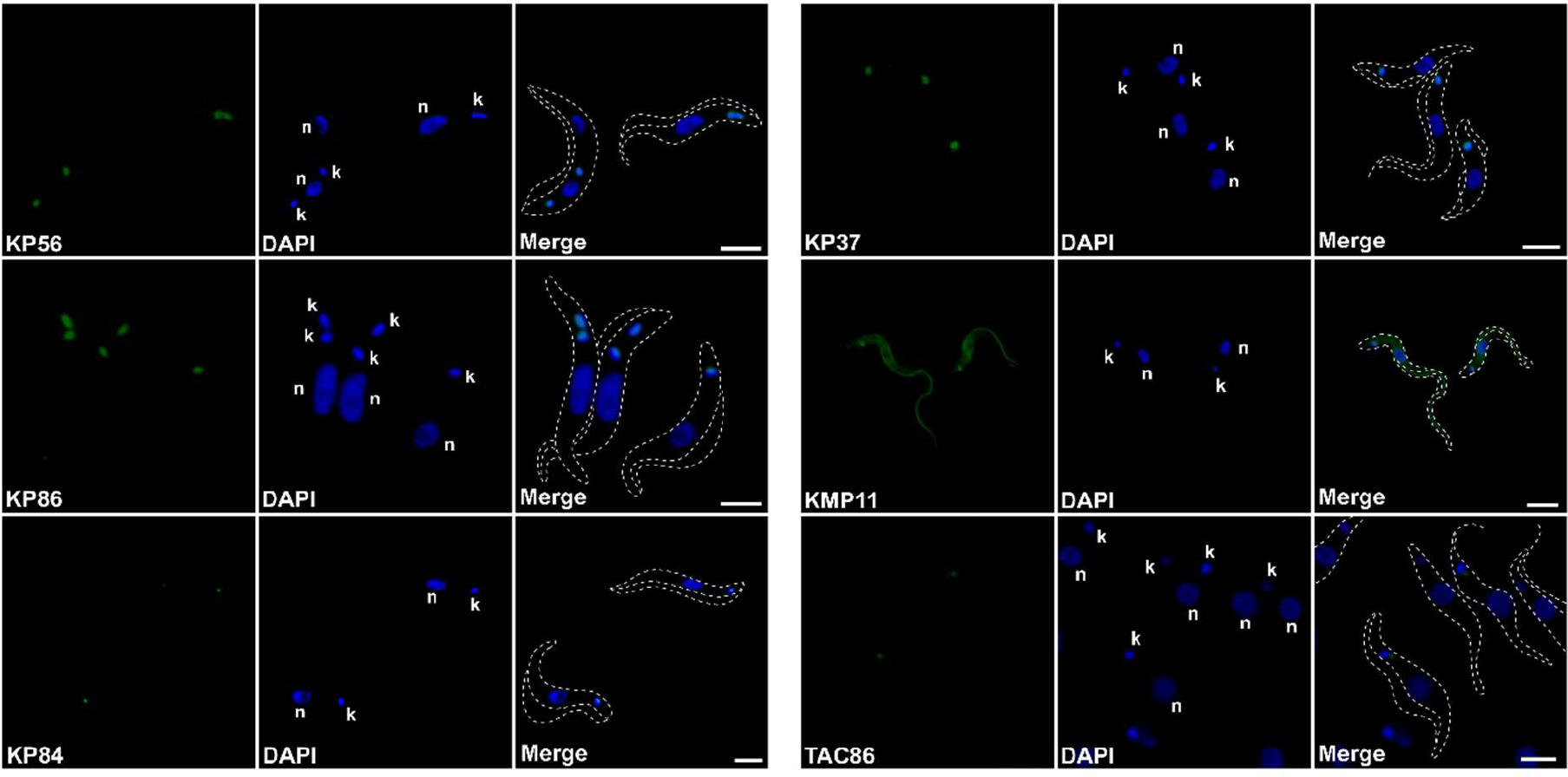
Localization of novel kinetoplast-associated proteins. Immunofluorescence assay of the candidate proteins reveals their localization within the kinetoplast. Nucleus (n) and kDNA (k) were stained with DAPI (blue) and the respective V5-tagged protein is shown in green. The occurrence of cyan shows co-localization of the protein in respect to or proximal to the kinetoplast. Scale bar = 2 μm.

Out of the initial candidates subjected to V5 tagging, one protein annotated as ‘HD domain containing protein’ (Tb927.7.4810) as well as ‘phenylalanyl-tRNA synthetase subunit alpha’ (Tb927.11.14120) were repeatedly resistant to epitope generation, while tagged ‘phenylalanyl- tRNA β-subunit’ (Tb927.11.2360) did not exhibit a KP signal and was predominantly enriched in the cytosolic fraction **(SFig. 1A, B)**. The remaining candidates were treated with digitonin to obtain crude cytoplasmic and mitochondrial fractions. Western blot analysis revealed mitochondrial localizations **(SFig. 2)** which was further confirmed by IFA **(Fig. 1)**.

**Figure 2.**
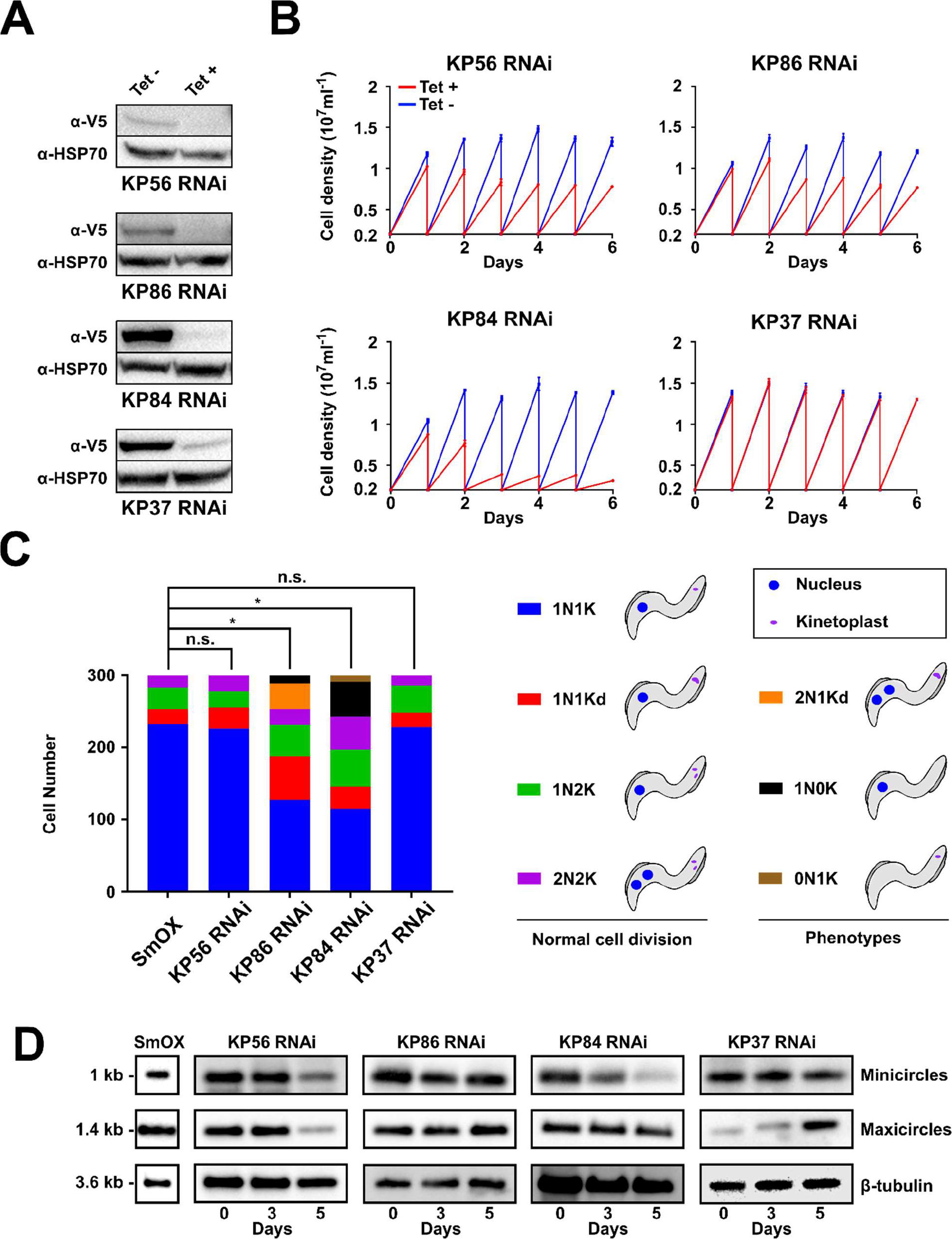
Effects of RNAi-based KPs depletion on cell viability, division and kDNA levels. (**A**) Western blot analysis of RNAi knock-down cells. The expression levels of V5-tagged proteins were measured 72 hours post-induction (hpi) with tetracycline and compared to the corresponding non-induced cell line. HSP70 protein served as a loading control. (**B**) Graphs depicting growth curves as cell numbers (10^7^ cells/ml) vs. days post-RNAi induction, in which cultures were diluted to 2x10^6^ cells/ml every 24 hours (n=3; error bars, SD). (**C**) Cell cycle analysis. Three hundred DAPI-stained cells, either 72 hpi post- RNAi induction or non-induced wild types (WT), were categorized according to both the presence and number of nuclei (N), kinetoplasts (K), and dividing kinetoplasts (Kd). Percentages corresponding to each sub-population are indicated. Statistically significant (*, p<0.05) differences were calculated by Student *t*- test paired contrasts with respect to the WT cells. Physiological and aberrant configurations of nuclei (N) and kinetoplasts (K) were quantified for each RNAi-induced cell line. Schematic view of division of the WT and RNAi-interfered cell lines, in which sequential steps of normal cell division are diagramed. (**D**) Southern blot analysis following kDNA-related phenotypes. Total genomic DNA was isolated from the WT cells, non-induced cells and those 3- and 5-days post-RNAi induction, resolved on agarose gels following digestion with restriction enzymes, blotted and hybridized with maxicircle- and minicircle- specific probes. Probes against cytosolic β-tubulin were employed as a loading DNA control. Sizes are indicated in kb.

New kinetoplast proteins were assigned the abbreviation of KP (<u>K</u>inetoplast <u>P</u>rotein) along with their molecular weight, generating a set composed of KP37 (Tb927.6.4510), KP56 (Tb927.8.4040), KP84 (Tb927.7.5320), and KP86 (Tb927.10.15660). Previously annotated KMP-11 (Tb927.9.13820), showed a signal corresponding to both the flagellum and the TAC region adjacent to the kinetoplast, similar to localizations reported previously (Z. Li & Wang, 2008).

A sixth tagged protein exhibited a faint signal proximal to the kinetoplast **(Fig. 3)**, likely due to its low endogenous expression or stability **(SFig. 3A, B)**. IFA difficulties were ultimately mitigated by the generation of an overexpressing cell line **(SFig. 3C)**, which displayed a clearly visible signal adjacent to the kinetoplast, but was accompanied by severe morphological phenotypes, such as bloated cells. Similar to KMP11, intracellular localization of this protein is immediately adjacent to the kinetoplast, indicating localization in the TAC filaments, for which we designate this protein TAC86. Thus, we initially established kinetoplast-associated localizations for six novel proteins **(Table 1)**.

**Figure 3.**
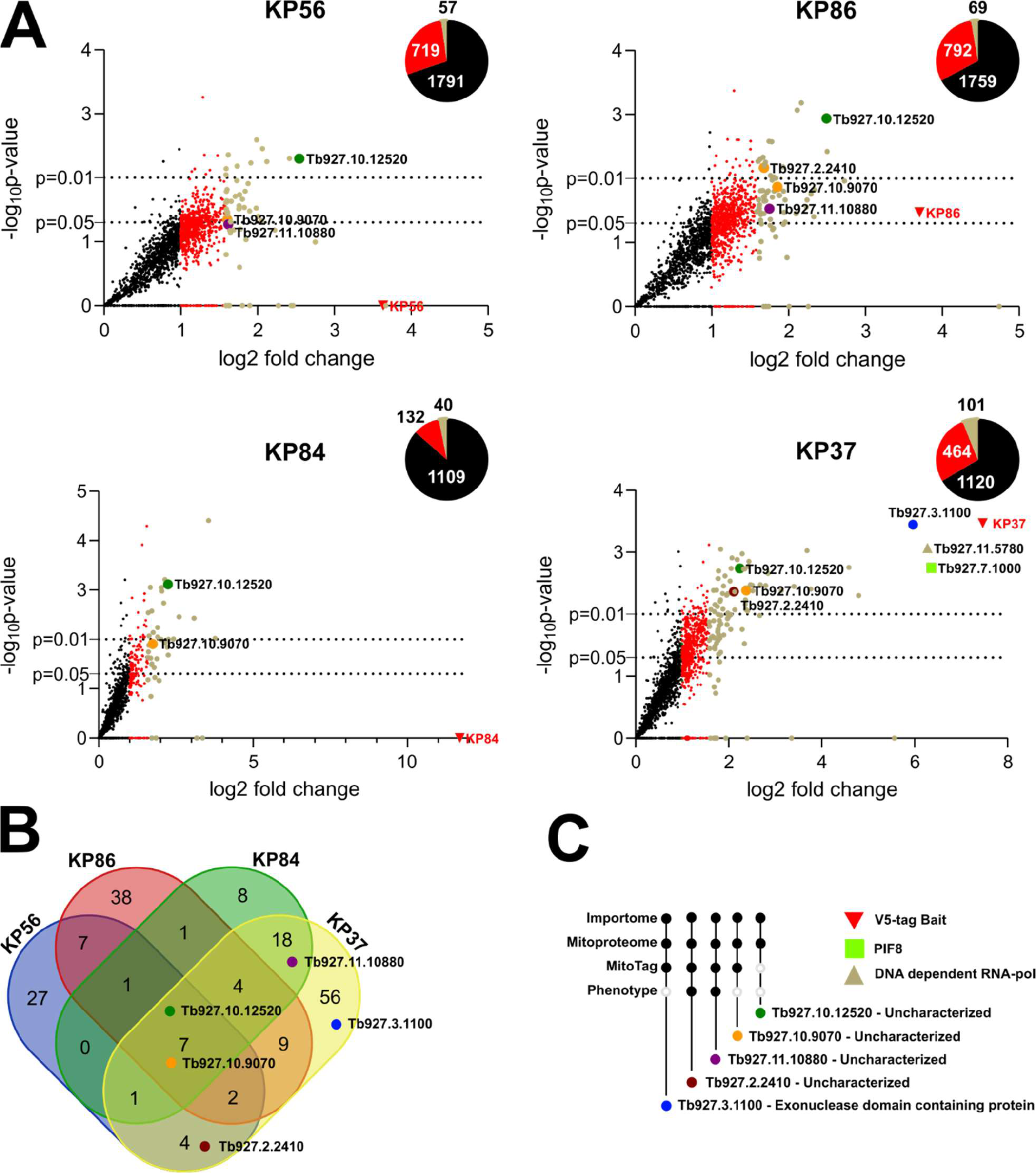
Putative interactors of kinetoplast-associated proteins and new putative kinetoplast proteins. (**A**) Volcano plots showing the results of 4 independent pulldown assays using V5-tagged KPs as bait, graphed as -log_10_ transformed p-values for each retrieved protein as a function of log_2_- transformed fold-enrichment as compared with untagged WT SmOx cells. Bait proteins (red triangles) and shared interactors of interest (colored circles) are indicated. Green dots correspond to other ≥ 3-fold enriched proteins, accessions are provided in **S** Table 1. Smaller red dots represent ≥2<3-fold enriched proteins and black dots are remnant enriched proteins. Pie charts refer to the total number of proteins recovered in immunoprecipitations versus the number of ≥ 3-fold enriched proteins. Significance thresholds of p=0.01 and p=0.05 p-values are indicated by dotted lines. (**B**) Comparative analysis of shared co- immunoprecipitating partners. Venn diagram showing conspicuous interactors present in one or more co- immunoprecipitation assays. (**C**) Prioritized proteins subjected to downstream validation. Protein presence in previous studies (Importome, Mitoproteome or MitoTag) or growth phenotype in RNAi screen also indicated.

**Table 1.**
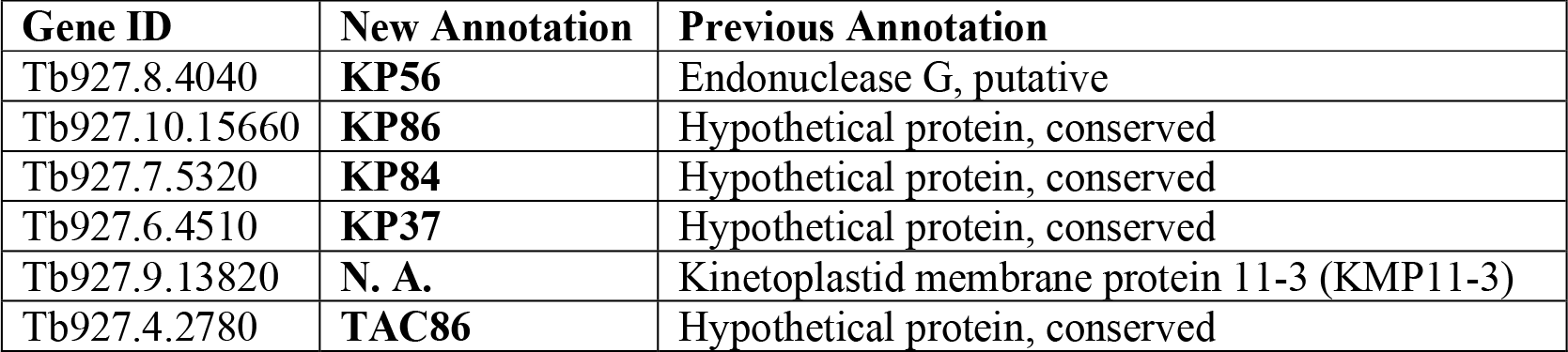
Nomenclature and localization of the novel kinetoplast-associated proteins. Gene identifiers on TriTrypDB.org with both new (from this study or Not Applicable [N. A.] if previously annotated manually) and previous annotations shown.

### KP56, 84 and 86 display growth phenotypes following RNAi

To examine whether the four novel kinetoplast proteins are required for viability, we generated inducible RNAi knock-down cell lines (RNAi KD) in the corresponding V5 tagged as parental genetic background. The full or partial ablation of the tagged protein was assessed by western blot analysis. With the exception of KP37, all tagged proteins were not detectable 72 hours post- RNAi induction (hpi) **(Fig. 2A)**. Growth curves revealed that the RNAi KD of KP56 and KP86 resulted in moderate growth retardation, while the depletion of KP84 resulted in a more drastic phenotype, with a severe growth arrest occurring at 48 hpi, whereas KP37 RNAi KD cells did not exhibit any growth phenotype for the observed time frame (**Fig. 2B**). Given that we were unable to completely eliminate this epitope-labelled protein by RNAi (**Fig. 2A**), it is plausible that this residual protein presence was sufficient to support growth in this cell line.

### KP84 and 86 RNAi lines exhibit aberrant cell cycles

To assess the cell cycle effects following the RNAi KD of kinetoplast-associated proteins, DAPI-stained cells were analyzed at 72 hpi and compared to non-induced cells. <u>W</u>ild type (WT) *Trypanosoma brucei* cells divide in a stepwise process in which the kinetoplast (K), nucleus (N) and single copy organelles, such as the endoplasmic reticulum, Golgi apparatus and flagellum, must duplicate, segregate and distribute between daughter cells in a highly regulated manner (Sherwin & Gull, 1989). Under physiological conditions, kinetoplast duplication in this species precedes nuclear division and cells progress from 1N1K to intermediate 1N1Kdividing(d), 1N2K, and 2N2K, reverting again to 1N1K after cytokinesis (Benz, Dondelinger, McKean, & Urbaniak, 2017) **(Fig. 2C)**.

For each cell line, 300 cells were examined and categorized into either physiological (1N1K, 1N1Kd, 1N2K, 2N2K) or aberrant phases (2N1Kd, 1N0K, 0N1K) **(Fig. 2C)**. The reduction of KP37 and KP56 resulted in a cell division profile significantly different from WT cells, with approximately 70% cells in G_1_/S 1N1K phase. The remaining 10%, 15% and 5% cells were distributed between the dividing stages categorized as 1N1Kd, 1N2K, and 2N2K, respectively.

Cells with downregulated KP86 exhibited a significant reduction of 1N1K cells (40% vs. 70%) and an increase of 1N1Kd (20% vs. 10%), as compared to the WT cells, suggesting arrested kinetoplast division **(Fig. 2C)**. Accordingly, 13% of aberrant 2N1Kd cells were observed, exhibiting fully divided nuclei but with unresolved kinetoplast division as well as 4% of 1N0K cells, in which the undivided kinetoplast was not inherited after cytokinesis.

For the KP84 RNAi KDs, which also displayed a gross effect on cellular division, significantly reduced numbers of 1N1K cells (35% vs. 70%) were accompanied with an increased number of 2N2K cells (18%). In this cell line we observed a higher proportion of 1N0K akinetoplastic cells (15%), together with 2% of anucleated cells (0N1K) **(Fig. 2C)**.

### kDNA abundance is altered in KP37, 56 and 84 RNAi

Kinetoplast-associated proteins, when subjected to RNAi KDs, knock-outs and overexpression, frequently trigger phenotypes affecting kDNA inheritance and alterations in kDNA content (Amodeo et al., 2022; Liu et al., 2009). To investigate this, we monitored the levels of maxicircles and minicircles relative abundance by southern blot analysis in RNAi KD cells **(Fig. 2D)**.

Indeed, in cells depleted for KP56, a parallel decrease of both maxicircles and minicircles occurred 5 days post-RNAi induction. Interestingly, no significant differences in cell cycle ratios were exhibited at 72 hpi, despite the appearance of reduced cell growth starting from 24 hpi, the latter phenotype attributable to the loss of maxicircles **(Figs. 2B, C)**. Equally, while KP86 RNAi KD showed significantly altered growth and cell cycle phenotypes, KP86 RNAi KD shows no changes in the maxicircle and minicircle signals **(Fig. 2D)**, suggesting a function not directly pertaining to kDNA replication. A clear impact on minicircles were observed in cells in which KP84 had been noticeable from 72 hpi. The extreme growth phenotype of KP84 **(Fig. 2B),** as well as the large proportion of aberrant dividing cells at 72 hpi **(Fig. 2C)** suggest a near-immediate phenotype, most likely a consequence from the extreme reduction of minicircles, affecting the successful editing of mitochondrial transcripts. A peculiar phenotype was noticed in KP37 RNAi cells. Despite the lack of phenotypes for cell growth **(Fig. 2A)** or cell cycle progression **(Fig. 2C)** within the observed time frames, KP37 RNAi KD nonetheless exhibits an accumulation of maxicircles at day 5, suggesting a negative regulatory role within maxicircle maintenance.

### Identification of shared proteins through KP immunoprecipitation and mass spectrometry

To determine putative-interaction partners of KP37, KP56, KP84, and KP86, each V5-tagged cell line was employed as bait for immunoprecipitation from the crude mitochondrial lysates of procyclic *T. brucei*. Pulldowns of each bait protein were analyzed by quantitative mass spectrometry and compared to WT cells **(Fig. 3A)**. Overall, a total of 209 unique interacting partners with significant (≥ 3-fold) enrichment were identified between the pulldowns obtained from these 5 cell lines.

Comparative analysis of the pulldown data revealed that although the examined tagged kinetoplast-associated proteins do not co-precipitate with each other, they pulled down several shared, potentially interacting partners (**Fig. 3A)**. From the 209 proteins aforementioned proteins, mitochondrial localization was predicted for 45 and 28 proteins by MitoFates (Fukasawa et al., 2015) and TargetP (Armenteros et al., 2019), respectively. Equally, their previously documented presence in the mitochondrion of *T. brucei* was referenced against the MitoTag database (108/209) (Pyrih et al., 2023), the mitoproteome (70/209) (A. K. Panigrahi et al., 2009) and the ATOM40 depletome (92/209) (Peikert et al., 2017).

A comprehensive analysis of mutual identified proteins protein revealed a group of 15 sequences either shared in all (seven) or the majority of immunoprecipitations (eight) **(Fig. 3B).** Of interest, the majority of these proteins either have no annotation, or exhibited homology-inferred annotations related to DNA or RNA metabolism **(Table 2).**

**Table 2.** Putative novel kinetoplast-associated proteins co-immunoprecipitated in the majority of analyzed KP pulldowns. Gene identifiers and annotation on TriTrypDB.org.

| Gene ID | Annotation |
| --- | --- |
| *Tb927.9.4290* | hypothetical protein, conserved |
| *Tb927.11.5330* | hypothetical protein, conserved |
| *Tb927.10.12520* | hypothetical protein, conserved |
| *Tb927.11.1100* | cysteine peptidase, Clan CA, family C2, putative |
| *Tb927.2.100* | retrotransposon hot spot protein 1 (RHS1), putative |
| *Tb927.10.9070* | hypothetical protein, conserved |
| *Tb927.8.5580* | N-terminal region of Chorein, a TM vesicle-mediated sorter/Integral peroxisomal membrane peroxin, putative |
| *Tb927.7.5320* | hypothetical protein, conserved |
| *Tb927.2.4710* | RNA-binding protein, putative |
| *Tb927.11.10880* | hypothetical protein, conserved |
| *Tb927.4.2080* | Coiled-coil and C2 domain-containing protein |
| *Tb927.7.2770* | hypothetical protein, conserved |
| *Tb927.6.1070* | hypothetical protein, conserved |
| *Tb927.7.2330* | Protein of unknown function (DUF1663), putative |
| *Tb927.3.4140* | hypothetical protein, conserved |

### Two novel Kinetoplast proteins identified from prioritized co-precipitation candidates

Based on these results, we endeavored to investigate a select group of these interacting proteins, to determine their localizations. We manually prioritized four proteins (Tb927.10.9070, Tb927.10.12520, Tb927.11.10880 and Tb927.2.2410) that were enriched in multiple pulldowns, were additionally present in two previous *T. brucei* mitochondrial proteomes (A. Panigrahi et al., 2009; Peikert et al., 2017). We additionally considered the annotation of candidates, choosing a fifth protein (Tb927.3.1100), based on the presence of an putative exonuclease domain, as well as whether any of these proteins appeared essential from the procyclic stage of *T. brucei* in a high-throughput RNAi analysis (Alsford et al., 2011) **(Fig. 3C)**.

The five selected proteins were endogenously tagged with a V5 epitope similar to other previous candidates, and their correct integration as well as mitochondrial localization were conducted as described above. Reassuringly, all prioritized candidates appeared in the mitochondrial fraction **(SFig. 4)**, while immunofluorescence analysis demonstrated an enriched kinetoplast signal for two proteins (Tb927.3.1100 and Tb927.10.9070) **(Fig. 4)**. Accordingly, we name these two new *bona fide* kinetoplast proteins as KP45 and KP66, respectively. These proteins appear unique in that they display both a kinetoplast signal as well as reticulated mitochondrial pattern, suggesting a dual localized signal shared with the mitochondrial matrix **(Fig. 4)**.

**Figure 4.**
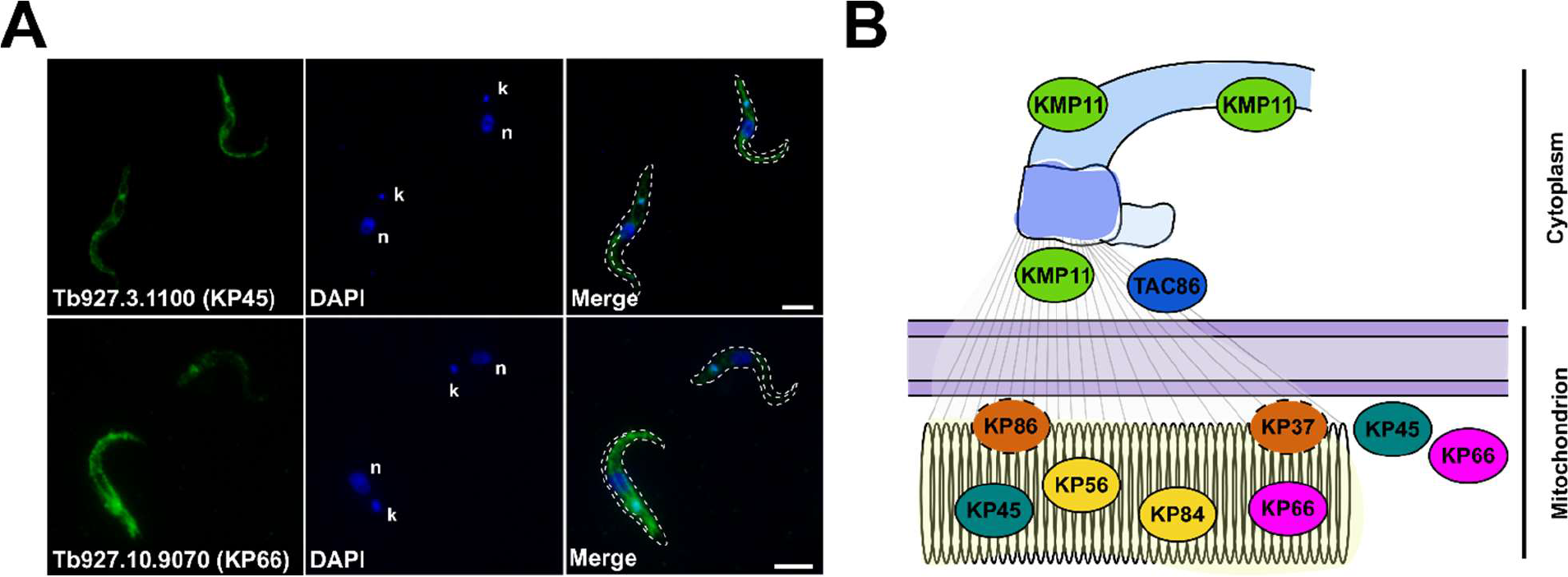
Novel kinetoplast-associated proteins identified in KP pulldowns. (**A**) Immunofluorescence analysis of KP45 and KP66. Nucleus (n) and kDNA (k) were stained with DAPI (blue) and the respective V5-tagged protein is shown in green. Scale bar = 2 μm. (**B**) Schematic representation of all kinetoplast-associated proteins characterized in this study. The validated, approximate topology of proteins within the TAC is also indicated.

### Phylogenetic distribution, conservation, and functional prediction of novel kinetoplast-associated proteins

Phylogenetic distributions of these eight novel kinetoplast-associated genes from *T. brucei* were surveyed against a spread of available kinetoplastid genomes as well as transcriptomes and genomes from Euglenozoan sister clades of euglenids and diplonemids. The more distantly related *Homo sapiens* and *Saccharomyces cerevisiae,* of the Opisthokont supergroup, were additionally included. These novel kinetoplast proteins showed varying levels of conservation and distribution, with rare occurrences of apparent gene loss amongst surveyed organisms **(Fig. 5A)**. Three genes are present outside the clade of kinetoplastids, and thus preceded the emergence of the kinetoplast itself. KP56, annotated as an endonuclease, shows the broadest distribution, being present in both *H. sapiens* and *S. cerevisiae*. KMP11 was likely present in the last Euglenozoan common ancestor, as seen from its presence in all three clades of Euglenozoa, despite also exhibiting the largest number of apparent losses from this surveyed selection. KP45 is confined to a smaller subgroup of Euglenozoa termed ‘glycomonada’, encompassing both diplonemids and kinetoplastids.

**Figure 5.**
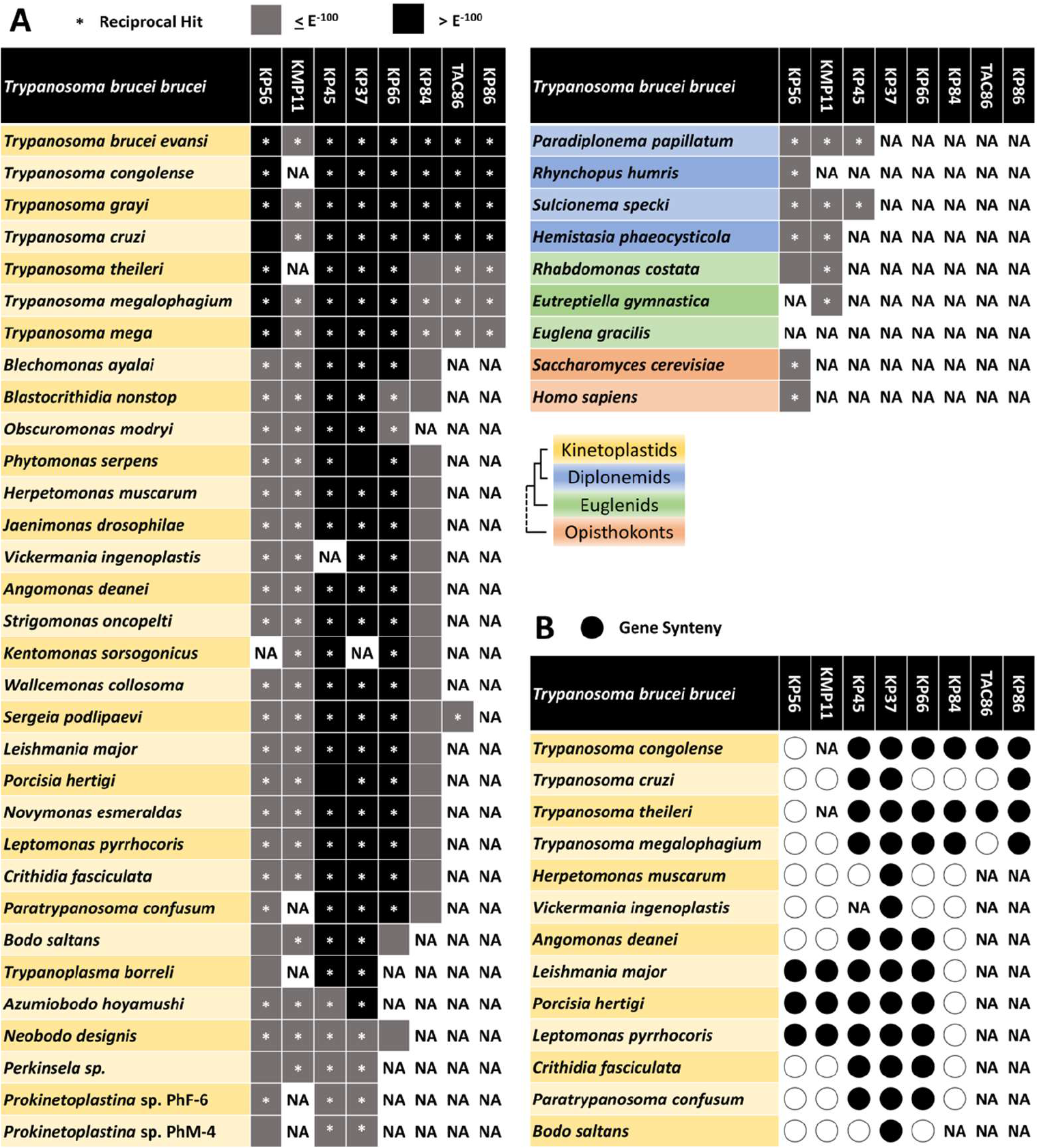
Conservation of kinetoplast-associated proteins within and outside of the phylum Euglenozoa. (**A**) The presence, score values and reciprocal BLAST hit retrievals for orthologs of all eight novel kinetoplast-associated proteins in members of the phyla Euglenozoa and Opisthokonta. Black and grey squares represent BLAST scores higher than E^-100^ and E^-50^, respectively. Gene absences represented by NA (<u>N</u>ot <u>A</u>pplicable). (**B**) Synteny of analyzed genes within the family Trypanosomatidae. Lack of gene synteny represented by empty circles. Gene accessions and score values in **S** Table 2.

Of the five genes observed only in kinetoplastids, two components (KP37 and KP66) are present in free living members such as *Neobodo designis* and *Bodo saltans*, while the remaining three (KP86, TAC86 and KP84) are found only in an exclusively parasitic clade **(Fig. 5A)**. KP86 appears to be the most recently developed gene, while the presence of TAC86 in the more distantly related *Sergeia podlipaevi*, suggests the loss or divergence beyond recognition of this gene amongst more closely related members to *T. brucei*. Interestingly, the dyskinetoplastic *T. brucei evansi,* despite notably lacking kDNA, nonetheless retains a full ‘complement’ of these novel genes, suggesting minimal time of divergence from the model organism *T. brucei brucei*.

The colinear order of genes between species, concerning their immediately adjacent genes, referred to as synteny, was additionally assessed across high quality genomes of kinetoplastids. Complete synteny was seen in all members possessing the most restricted gene of KP86, as well as the more broadly distributed KP37 **(Fig. 5B)**. KP37 is the only syntenic gene amongst both *Herpetomonas muscarum* and *Vickermania ingenoplastis*, who otherwise appear to have undergone extensive levels of genome rearrangement. Furthermore, KP37’s high sequence conservation and reciprocal hit retrieval rate suggest a high degree of essentiality for this gene, both in terms of its conservation as well as its genomic position **(Fig. 5A&B)**, in spite of no immediate growth or cell cycle phenotype observed from knockdown studies employed here **(Fig. 2)**. Both KP45 and KP66, identified via immunoprecipitation in this study, additionally show similar levels of synteny, conservation and distribution among kinetoplastids to KP37 **(Fig. 5A&B)**, prompting speculation on the precise molecular functions these genes encode for.

## DISCUSSION

An unexpected finding of the MitoTag study was the demonstration that the mNeonGreen tag and other large globular epitopes produced a localization proximal to the kinetoplast, affecting mitochondrial proteins which would normally sub-localize in the matrix or integral inner membrane, a localization termed kinetoplast proximal enrichment. This specific localization is similar, but ultimately distinct from the signal elicited by many genuine kinetoplast proteins tagged in a similar manner (Pyrih et al., 2023). In turn, this phenomenon enabled the identification of several new kinetoplast candidates from the MitoTag dataset. Despite this, such a similarity between the genuine and artificial kinetoplast signals naturally warrants skepticism of any new proteins identified as ‘kinetoplast’ *via* this tagging strategy. Thus, it was viewed as a necessity to employ the smaller, non-globular V5 tag, which specifically does not induce such artificial interactions with the kinetoplast (Pyrih et al., 2023). Accordingly, if these proteins constitute genuine components of the kinetoplast, then a kinetoplast signal should persist with this epitope. In summary, of the 14 candidates that were ultimately chosen for further analysis, eight were validated as genuine kinetoplast-associated components through IFA.

To the best of our knowledge, 57 *T. brucei* proteins have so far been associated either by location and/or function with the kinetoplast (Hoffmann et al., 2018; Jensen & Englund, 2012; Lindsay et al., 2008; Liu et al., 2009; Povelones, 2014), and the discovery and validation of eight novel proteins belonging to this structure therefore represents a significant development for a hallmark structure of *T. brucei*, arguably the best studied protist within the eukaryote domain (Billington et al., 2023; Lukeš et al., 2023; Marques, Ridgway, Tinti, Cassidy, & Horn, 2022; Pyrih et al., 2023).

From the recent Hyper-LOPIT spatial resolution of *T. brucei* (Moloney et al., 2023), our novel kinetoplast proteins are listed simply as constituents of the mitochondrial matrix. While this study achieved excellent resolution of mitochondrial compartments and complexes, including a sub- fraction of the matrix, enriched in ribosomal subunits, the kinetoplast was not resolved in a similar manner. We suggest that difficulties in defining the kinetoplast through interaction and fractionation-dependent assays stem from the fragile nature of this complex, with cell disruption releasing most kinetoplast components freely into the matrix. As a result, tagging studies remain the ideal method for identifying and characterizing future components of the kinetoplast.

The recent availability of many genome and transcriptome assemblies, covering the diversity of Euglenozoa, and Trypanosomatidae in particular (Albanaz et al., 2023; Alexei Yu Kostygov et al., 2024), prompted us to investigate the distribution of orthologs, revealing markedly varying patterns of distribution, with three genes (KP56, KMP-11 and KP45) preceding the development of the kinetoplast, likely being recruited and functionally adapted to this complex after its emergence.

KP56, of particular interest, is distributed beyond Euglenozoa, including orthologs in yeast and humans, which represents the predominant endo/exonuclease embedded within the inner mitochondrial membrane, performing functional roles in apoptosis initiation as well as mitochondrial DNA repair and recombination (Cymerman, Chung, Beckmann, Bujnicki, & Meiss, 2008; Rosamond, 1981). Like its mammalian counterpart, which has undergone an ancient duplication and subsequent functional specializations, a more recent duplication in *T. brucei* has resulted in both Tb927.8.4090 and KP56 (Tb927.8.4040), which are distinguished by a single nucleotide in their ORFs (translating to an Alanine to Valine substitution between KP56 and Tb927.8.4090 respectively) (Gannavaram, Vedvyas, & Debrabant, 2008). While Tb927.8.4090 was not targeted in this study (**Suppl Fig. 5**) and exhibits inconclusive localization from TrypTag (Billington et al., 2023), we expect this protein to also be localized to the kinetoplast due to its extreme similarity to KP56. Importantly, our RNAi results show a distinct growth phenotype in KP56, accompanied by a reduction in both maxicircles and minicircles, which stands in contrast to previous RNAi work reporting no effect on growth, likely due to an inability of this previous study to completely eliminate targeted mRNA (Gannavaram et al., 2008).

KMP11 has been experimentally characterized as a conserved and broadly distributed protein within the Trypanosomatidae lineage, of which knock-down studies affect cytokinesis and formation of the flagellar attachment zone (Diez et al., 2008; Z. Li & Wang, 2008). KMP11 has previously been referred to as a marker for the flagellum and the basal body (Z. Li & Wang, 2008). However, MitoTag, using signal spacing relative to the kinetoplast, positioned this protein with other TAC proteins, immediately proximal to the basal body (Pyrih et al., 2023). Similarly, in our study, KMP11 shows a signal both adjacent to the kinetoplast and throughout the flagellum, that is ultimately never connected. Therefore, we argue that this protein should more properly be referred to as a constituent of TAC filaments, as opposed to the basal body.

While KP45 possesses an exonuclease domain and has been recorded in multiple mitochondrial investigations in *T. brucei*, this protein remains functionally uncharacterized. Its exclusive recovery from KP37’s immunoprecipitation, along with TbPIF8 helicase and a DNA-dependent RNA polymerase suggest roles in kDNA maintenance (J. Wang, Englund, & Jensen, 2012).

KP45’s shared distribution with diplonemids is notable, considering that diplonemids have equally undergone extensive mitochondrial DNA expansion (Burger & Valach, 2018). While not concentrating and intercatenating mitochondrial DNA in a structure homologous to the kinetoplast, its expanded, repertoire of maxicircles and minicircles nonetheless would have likely required additional components associated with nucleic acid maintenance.

The remaining five kinetoplastid-specific proteins (KP37, KP66, KP84, TAC86, KP86) represent putative innovations associated with the emergence of the highly evolutionary and ecologically successful kinetoplastid parasites, as well as potential therapeutic targets for intervention against its multiple disease-causing members.

KP37’s putative role as a negative regulator of maxicircle quantity represents an intriguing mechanism of maintenance, not reported from any other kinetoplast protein. RNAi of TbPIF2 also affects maxicircles exclusively, but, by contrast, results in their rapid depletion (Liu et al., 2009). KP37’s distribution suggests an emergence in tandem with or immediately succeeding the kinetoplast itself, and thus may represent a component critical to its initial genesis. The ineffective generation of a complete RNA interference ablation for this protein further strengthens its indispensability. The endosymbiont-bearing *Kentomonas sorsogonicus* represents the only surveyed kinetoplastid to have lost this gene, also lacking KP56, and may offer insight on a diverged system of kDNA maintenance (Votýpka et al., 2014). In *T. brucei* specifically, we hope to complement our results obtained from KP37 via an overexpressing cell line to clarify the regulatory role performed by this protein.

KP66, in addition to KP45, exhibits dual-mitochondrial localizations, comprising both the kinetoplast and the mitochondrial lumen. Other kinetoplast-associated proteins also display dual mitochondrial localizations, such as pATOM36, present in the TAC filaments as well as the outer mitochondrial membrane, participating in protein import and translocation (Käser et al., 2016; Pusnik et al., 2012). Like pATOM36, these proteins may be functionally required beyond the kinetoplast within the mitochondrial matrix. Their status as soluble mitochondrial matrix proteins offers explanation as to why they were not identified as kinetoplast candidates through MitoTAG (Pyrih et al., 2023) (additionally exhibiting KPE as a matrix constituents), instead requiring identification in this study through co-immunoprecipitations. We restricted our co-immunoprecipitation follow-up screening to five candidates, resulting in these two additional successful identifications, but similar co-enriched candidates (see Table 2), many of which lack any annotation, remain uninvestigated.

KP84, along with KP56, do not possess a predicted mitochondrial N-terminal target peptide, which the majority of mitochondrial proteins rely on for import into this organelle (Desy, Schneider, & Mani, 2012). Their previous tagging resulted in associations with the kinetoplast only when an epitope was present at the N-terminus, thus leaving the C-terminus exposed (Pyrih et al., 2023), which we replicated here. These traits are shared with another characteristic kinetoplast protein, DNA topoisomerase II (Tb927.9.5590), serving as a prototypical cell-cycle dependent marker of the antipodal sites (Amodeo et al., 2022), and additionally being involved in minicircle maintenance like the former two proteins (Z. Wang & Englund, 2001). Given the striking rarity of mitochondrial proteins dependent on their C-termini for organellar import, the presence of three kinetoplast proteins with such a dependency provides support for the existence of a dedicated trafficking pathway in *T. brucei* and other kinetoplastid flagellates for at least some kinetoplast-associated proteins, yet no other experimental data for such a mechanism are currently available.

TAC86’s localization, distinctly adjacent from the kDNA led us to designate this protein as a constituent of the TAC filaments, accordingly, this protein represents the only novel component not to be recovered from any previous mitochondrial analysis. There currently exist several monoclonal antibodies (BBA4/Mab22) that localize to the exclusion zone filaments of the TAC structure, for which protein counterparts in *T. brucei* are yet to be discovered (Schneider & Ochsenreiter, 2018). Therefore, TAC86 prompts further biochemical analysis to ascertain its exact functional and structural roles, as well as to determine whether it represents a previous unidentified antibody.

KP86 represents the most recent kinetoplast innovation identified in this study, being confined to the genus Trypanosoma. In *T. cruzi*, an ortholog to KP86 has been identified as an interacting partner of chromatin remodeling imitation switch ATPase (Díaz-Olmos, Batista, Ludwig, & Marchini, 2020). The abundance of aberrant RNAi cells exhibiting two nuclei and a dividing kinetoplast, accompanied by no reduction in probed maxicircles or minicircles, resembles the phenotype observed in a recently characterized segregation factor TbNAB70 (Tb927.11.7580) (Cadena et al., 2024). In turn, this suggests that KP86 fulfills a role in kinetoplast segregation, with is ablation leading to replicated kDNA that is unable to successfully divide, arresting the cell in a 2N1kdiv phenotype. KP86 thus represents the second essential identified factor affecting the segregation of kDNA. When compared directly, TbNAB70 RNAi cells exhibit a more extreme and immediate growth phenotype to KP86. While TbNAB70 ablated cells also exhibit a combination of aberrant 2N1K and 2N1Kdiv, with the proportion of dividing kinetoplasts decreasing over time, KP86 ablation was only observed producing 2N1Kdiv cells. Furthermore, endogenously tagged KP86 does not appear to localize to the filamentous connecting structure referred to as the ‘nabelschur’, suggesting different timings and mechanisms of action involved in coordinating successful segregation.

In conclusion, the work presented here has identified several novel proteins associated with the kinetoplast through both localization studies and functional characterization. The proteins identified here are of relevance to our understanding of core biological processes of cell division as well as mitochondrial DNA maintenance, and furthermore represent promising targets for drug development amongst the many kinetoplastid parasites in which these components are distributed. Our study encompasses the methodological approaches necessary to determine any remaining components of the kinetoplast, to conclusively resolve our understanding of this highly complex cellular structure.

## MATERIALS AND METHODS

### Generation of *T. brucei* transgenic cell lines

*T. brucei* SmOx cell line (Poon, Peacock, Gibson, Gull, & Kelly, 2012) procyclic forms were transfected by electroporation using the Amaxa apparatus, set with program X-14. After transfection, cells were seeded in 24-well plates in serial dilutions and selected by the appropriate resistance marker. For endogenous V5-tagging cell line generation, a previously described strategy was employed (Durante et al., 2020). N- or C-terminal endogenous tagging amplicons were obtained using long primers targeting a modified pPOTv4 vector containing a 3xV5 epitope (Dean et al., 2015). Cells were selected with Hygromycin 50 µg/ml and screened for 3xV5 expression by WB.

Given the extreme similarity between targeted gene Tb927.8.4040 (KP56) to Tb927.8.4090, additional steps were taken to ensure correct integration. Nuclear DNA of this transfected cell line was isolated via a GeneAll genomic extraction kit and underwent PCR amplification of the approximately 500bp region upstream of the newly integrated DNA, encompassing and extending beyond the native 5’ UTR to the point of nucleotide differentiation between these two genes. The amplified region underwent sanger sequencing through Eurofins Genomics and was aligned against upstream regions of Tb927.8.4040 and Tb927.8.4090 to ensure correct integration had taken place (**Suppl Fig 5**).

For generation of RNAi cell lines, PCR products targeting a specific optimal CDS regions minimizing off-target effects as determined by RNAit (Redmond, Vadivelu, & Field, 2003) were cloned BamHI/HindIII into the p2T7-177 vector. NotI linearized constructs were electroporated into the correspondent parental 3xV5 tagged cell line. Cells were selected with Phleomycin 2.5 µg/ml and the mRNA and protein KD were monitored by WB for against 3xV5 epitope, respectively. Oligonucleotides used for PCR amplification and acceptor plasmids are given in **Table S 3**.

### Immunofluorescence assay

For immunofluorescence assays, 10^6^ *T. brucei* cells were fixed with 4% paraformaldehyde for 30 min at RT. The cells were subsequently centrifugated at 1,300 x g for 5 min and resuspended in 200 μl PBS and placed on a glass slide for 30 min. Attached cells were washed once with PBS and fixed and permeabilized in methanol for 30 min at -20°C. The slides were washed three times with PBS and blocked with 5% milk inside a humid chamber for 1 h before being incubated with anti-V5 mouse primary antibody (Invitrogen) (1:400 dilution) overnight at 4°C. After three times wash with PBS, cells were incubated with secondary Alexa Fluor 488 goat anti-mouse IgG antibody (1:1000) (Invitrogen) for 1 h at RT. Slides were again washed three times with PBS before a final wash with ddH_2_O, air-dried, and mounted in ProLong Gold antifade reagent DAPI (Life Technologies). Images were captured with the BX51 Olympus Fluorescence Microscope and analyzed by ImageJ (Schindelin et al., 2012).

### Cytoplasm-mitochondrial fractionation and Western-blot analysis

Digitonin preparation of crude mitochondria was done according to an established protocol (Wenger, Oeljeklaus, Warscheid, Schneider, & Harsman, 2017). Briefly, to obtain crude mitochondrial fractions, 10^8^ *T. brucei* 3xV5 tagged cells were treated with 0.015% digitonin (w/v) on ice for 5 min. A fraction of this sample was collected as whole cell lysate (WCL). Mitochondrial concentrated fractions (mito) were obtained by centrifugation at 6,800 x g for 3 min at 4°C. The supernatant of this centrifugation was kept as the cytoplasmic fraction (cyto). All fractions were loaded and subjected to SDS-PAGE, transferred to PVDF membranes and analyzed by Western-blot (WB). Membranes were probed with primary 1:2,000 mouse anti-V5 (Sigma), 1:10,000 mouse anti-enolase, as control for cytoplasmic fractions, 1:10,000 mouse anti- HSP70 monoclonal antibody as a mitochondrial control, and secondary 1:2000 goat anti-mouse-HRP antibody (Sigma). After development with ECL chemiluminescence kit (Pierce), membranes were visualized and photographed on a Chemidoc (BioRad) analyzer.

### Growth curves and cell cycle analysis

For cell growth analysis, cultures were grown in triplicate in the presence and absence of the tetracycline-class antibiotic doxycycline for RNAi-induction at a final concentration of 1µg/ml. Cell concentrations (10^6^/ml) were measured using the Beckman Coulter Z2 Cell and particle Counter every 24 h and subsequently diluted to 2 x 10^6^ cells/ml, maintaining cells in exponential phase of growth. For the cell cycle analysis, uninduced WT and induced KD cell lines were mounted on microscopy slides, stained with DAPI and visualized under fluorescence microscope. Three hundred cells were categorized according to their nuclei and kinetoplast numbers.

Statistical analysis was performed by paired contrasts using the Student’s *t*-test implemented in the GraphPad software.

### Southern-blot quantification of maxi- and minicircles abundance

Total DNA was isolated from mid-log phase cultures before RNAi induction and on 3 and 5 dpi of RNAi. The cell pellets were washed in PBS and resuspended in TRI Reagent (Molecular Research Center) according to manufacturer’s guidelines. 5 μg of total DNA per lane were digested with HindIII and XbaI for tubulin and maxicircles-specific probes or RsaI in the case of minicircles-specific probes. Digested DNA was resolved on 0.75 % agarose gel and transferred to nylon membrane (Amersham). The hybridization was performed overnight using DIG Easy Hyb (Roche). Probes to detect tubulin, maxi- and minicircles were PCR amplified and labelled using PCR DIG Probe Synthesis Kit (Roche). The detection was done using DIG Luminiscent Detection Kit (Roche) following the recommendations of the manufacturer. Sequences of probes employed are included in **Table S3**.

### Immunoprecipitations

Immunoprecipitations (IPs) of 3xV5 tagged proteins was adapted from previously reported protocols (Cadena, Gahura, Panicucci, Zíková, & Hashimi, 2021). In brief, concentrated mitochondria obtained from crude mitochondrial isolates from 10^8^ cells were solubilized in IPP50 (50 mM KCl, 20 mM Tris-HCl; pH:7.7, 3 mM MgCl_2_, 10% glycerol, 1mM phenyl- methyl-sulfonyl-fluoride (PMSF) serine protease inhibitor, complete EDTA-free protease inhibitor cocktail (Roche) supplemented with 1.5% (v/v) digitonin for 1 h on ice. After centrifugation (18,600 x g for 20 min at 4°C), the supernatant was added to 1.5 mg of anti-V5- conjugated magnetic beads (Chromotek). The solubilized mitochondria were incubated under rotation with the beads for 90 min at 4°C. After the removal of the flowthrough, the beads were washed three times in IPP50 plus 1.5% digitonin. Prior to elution, the beads were transferred into a new tube. Elution was done in 0.1 M glycine; pH:2.0 for 10 min at 70°C with shaking at 1,000 rpm. The eluate was immediately neutralized with 1 M Tris; pH:8.0. The elution step was repeated once to achieve higher recovery. The input, eluate and flowthrough fractions were resolved by SDS-PAGE and analyzed by WB prior to LC-MS/MS analysis. All IPs were performed in a total of three biological replicates for statistical analysis.

### Mass spectrometry and analysis

Eluates of co-IP proteins were processed for LC-MS/MS analysis performed at Laboratory of Mass Spectrometry, Biocev, Charles University Faculty of Science facility and according to previously described protocols (Dibus, Korinek, & Cermak, 2022).

### Conservation and synteny survey of Kinetoplast proteins

Reciprocal BLAST searches were conducted against 8 Kinetoplast-associated genes against a variety of Kinetoplastid, Diplonemid and Euglenid and Opisthokont CDS, with a threshold E^-5^ value. Gene synteny was surveyed within genomes of Trypanosmatidae with N50 of 500kb or greater, in addition to *Paratrypanosoma confusm* and *Bodo saltans* (Albanaz et al., 2023).

### Data availability

The mass spectroscopy data have been deposited to the ProteomeXchange Consortium (http://www.proteomexchange.org) via the PRIDE partner repository with the data set identifier XXX.

## Acknowledgements

We thank Karel Harant and Pavel Talacko (Laboratory of Mass Spectrometry, Biocev, Charles University Faculty of Science) for performing LC-MS/MS analysis. We acknowledge support from the Czech Science Foundation (grants 22-01026S and 21-09283S to J.L.) and EU’s Operational Program “Just Transition” (grant CZ.10.03.01/00/22_003/0000003 LERCO to V.Y.), Ministry of Education, Youth and Sports of the Czech Republic (INTER-EXCELLENCE- LUASK22033 to V.Y) and Slovak Research and Development Agency (SK-CZ-RD-21-0038 to I. Š.-S.].

## Author Contributions

Conceptualization, I.M.D., M.H., L.R.C., and J.L.; methodology, M.S., L.R.C., C.B., V.R., I.Š.-S., M.H., and I.M.D.; investigation, M.S., L.R.C., C.B., V.R., I.Š., L.C., M.H., and I.M.D; data curation, L.R.C., M.H., V.Y. and I.M.D.; data analysis, L.R.C., M.H., and I.M.D.; writing – original draft, L.R.C., M.H., and I.M.D.; review and editing, all authors; visualization, I.M.D., M.H., L.R.C., and J.L.; supervision, I.M.D., V.Y. and M.H.; funding acquisition, J.L. and V.Y.

## Declaration of interests

The authors declare no competing interests.

## SUPPLEMENTARY DATA

**Supplementary** Table 1. Mass spectrometry data.

**Supplementary** Table 2. Orthologous genes analysis.

**Supplementary** Table 3. Primers and probes used in this study.

**Supplementary Figure 1.**
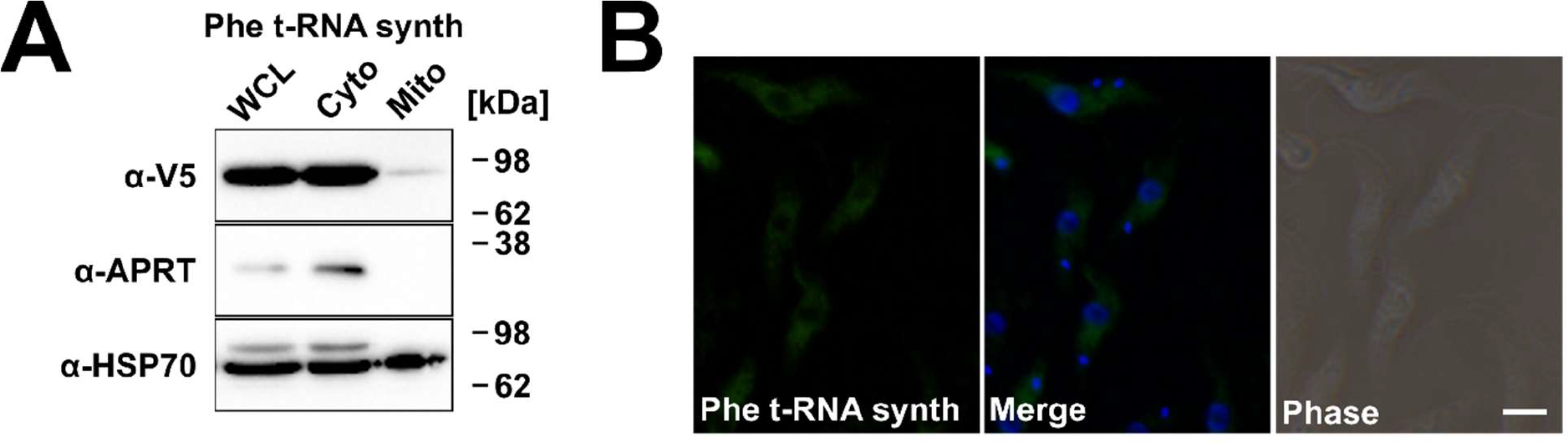
Localization of phenylalanyl-tRNA β-subunit (Tb927.11.2360). (**A**) WB analysis of crude cytoplasm-mitochondrial digitonin fractionations. Endogenously V5 tagged cells were subjected to crude digitonin fractionations and the distribution of V5 signal was assessed by WB with α- V5 antibody. Adenine phosphoribosyltransferase (APRT) and HSP-70 disrtibutions are shown as cytoplasm and mitochondrial controls, respectively. Molecular weights are indicated in kDa. (B) Immuno- fluorescence assay showing Tb927.11.2360 cytoplasmic localization. Nucleus (n) and kDNA (k) were stained with DAPI (blue) and the respective V5-tagged protein is shown in green. Scale bar = 2 μm.

**Supplementary Figure 2.**
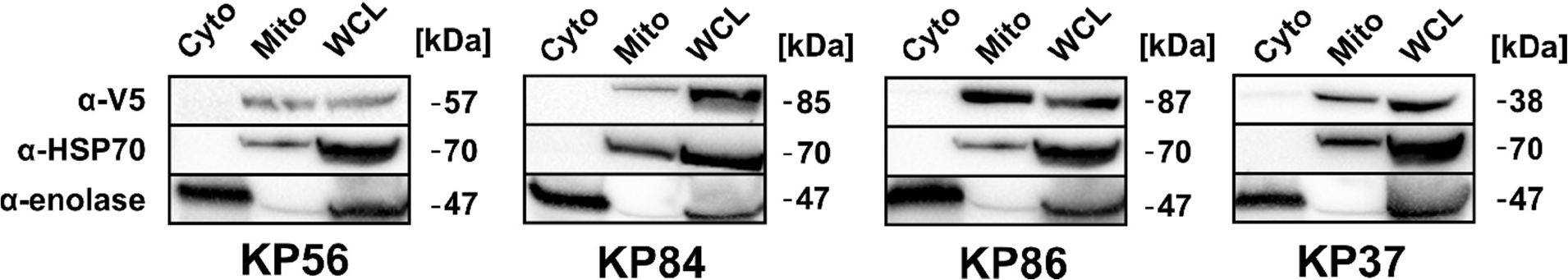
Subcellular localization WB analysis for novel kinetoplast proteins. (**A**) WB analysis of crude cytoplasm-mitochondrial digitonin fractionations. Endogenously V5 tagged cells were subjected to crude digitonin fractionations and the distribution of V5 signal was assessed by WB with α-V5 antibody. Enolase and HSP-70 distributions are shown as cytoplasm and mitochindrial controls, respectively. Molecular weights are indicated in kDa.

**Supplementary Figure 3.**
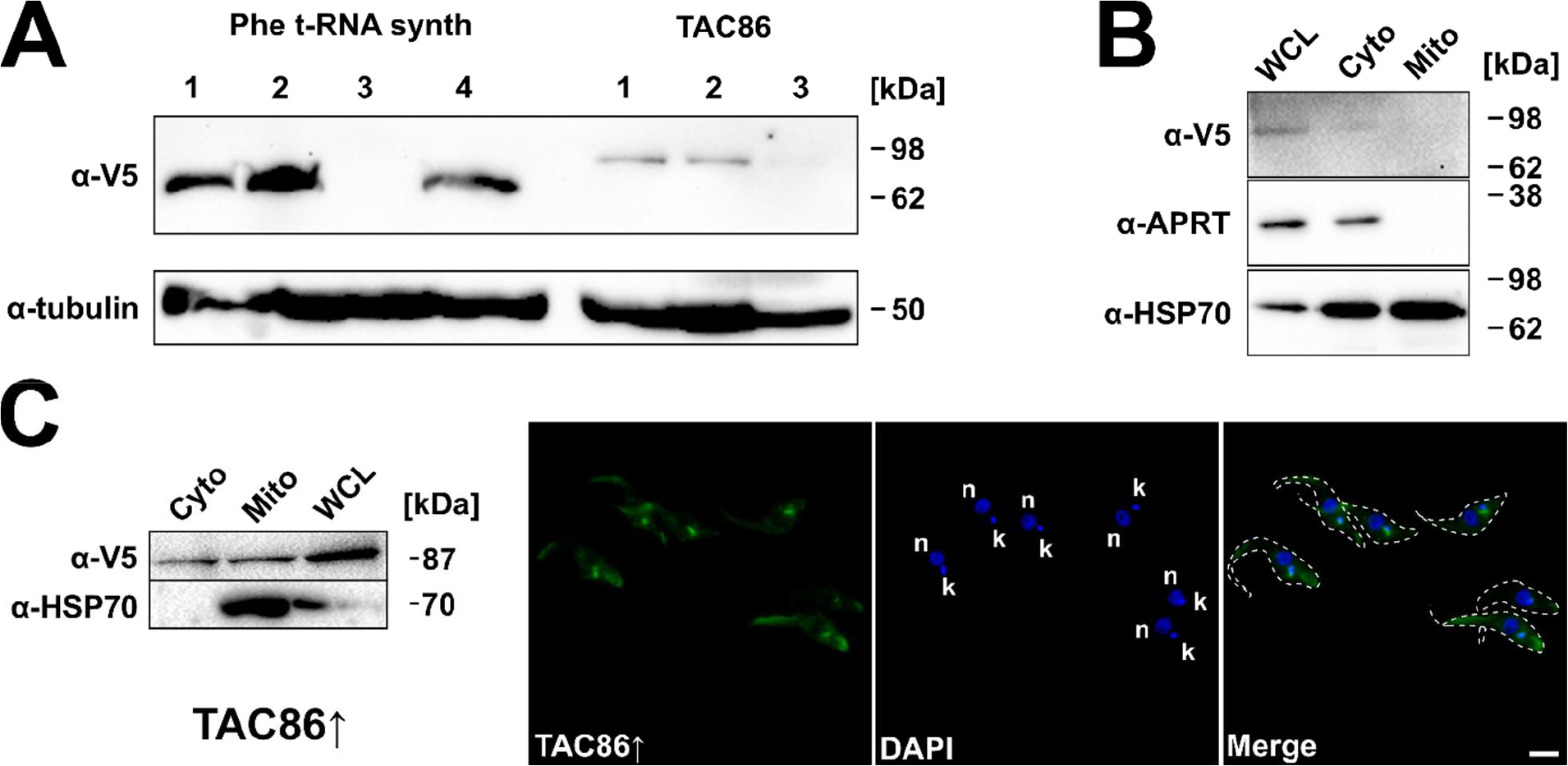
TAC86. (**A**) WB analysis of TAC86 relative endogenous expression. Clones of endogenously tagged TAC86 were compared in terms of relative WB signal over total lysates with expression of both Tb927.11.2360 and tubulin. (**B**) TAC86 fractionation over crude cytoplasm- mitochondrial digitonin fractionations. Endogenously V5 tagged cells were subjected to crude digitonin fractionations and the distribution of V5 signal was assessed by WB with α-V5 antibody. APRT and HSP- 70 disrtibutions are shown as cytoplasm and mitochindrial controls, respectively. Molecular weights are indicated in kDa. (C) TAC86-overexpressing cell line. Expression of TAC86 in induced cells was evaluated. TAC86 was detected both in mitochondrial and citoplasmic fractions. (C) Immuno- fluorescence assay showing TAC86 cytoplasmic and kinetoplast-proximal localization. Nucleus (n) and kDNA (k) were stained with DAPI (blue) and the respective V5-tagged protein is shown in green. Scale bar = 2 μm.

**Supplementary Figure 4.**
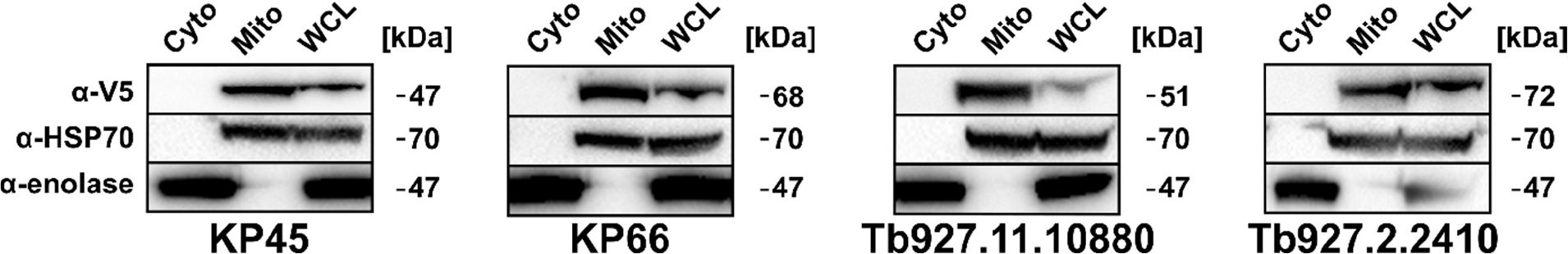
Subcellular localization WB analysis for pull down retrieved kinetoplast proteins interactors. (**A**) WB analysis of crude cytoplasm-mitochondrial digitonin fractionations. Endogenously V5 tagged cells were subjected to crude digitonin fractionations and the distribution of V5 signal was assessed by WB with α-V5 antibody. Enolase and HSP-70 disrtibutions are shown as cytoplasm and mitochindrial controls, respectively. Molecular weights are indicated in kDa.

**Supplementary Figure 5.**
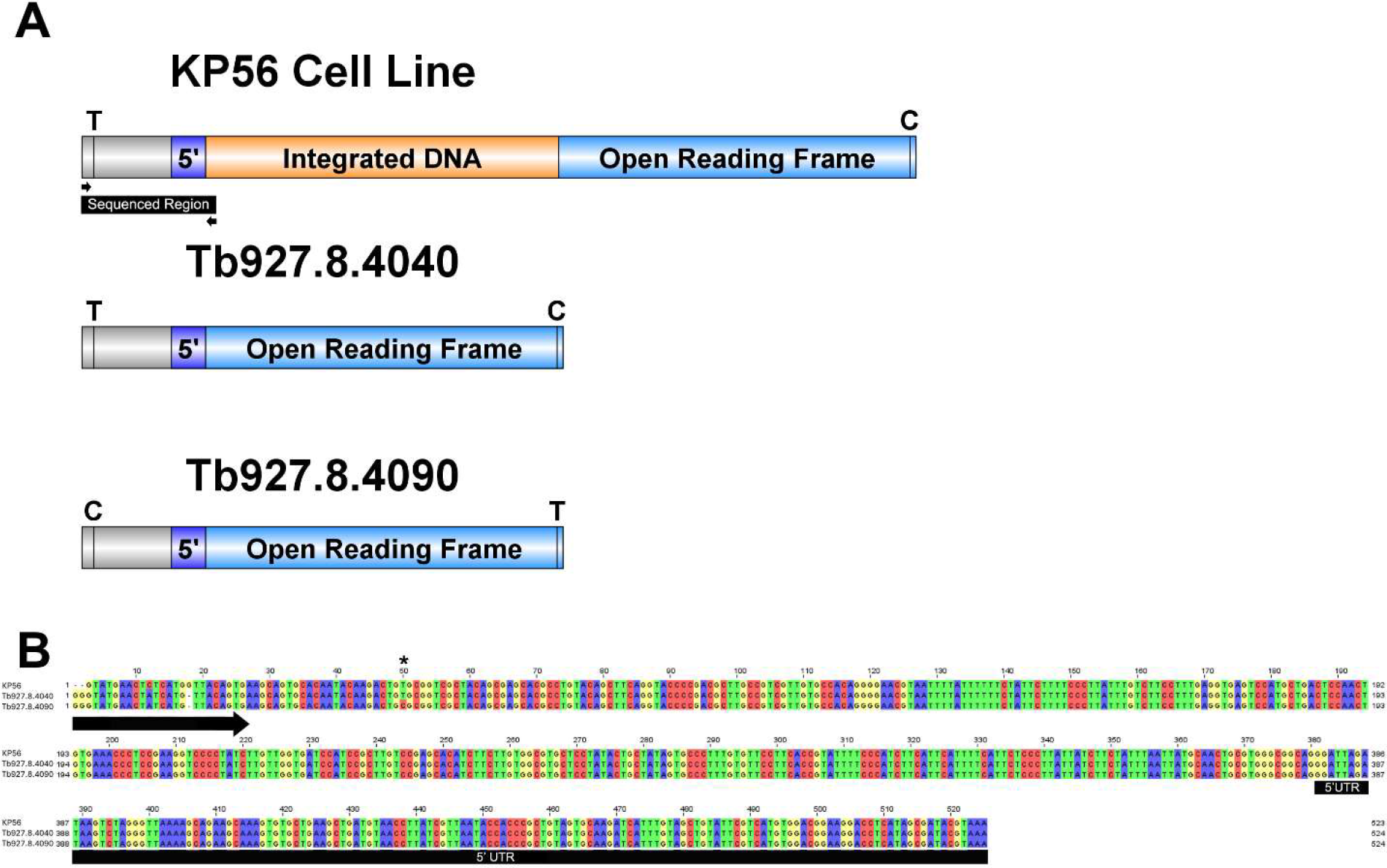
A. Schematic depiction of KP56 cell line as well as Tb927.8.4040 and Tb927.8.4090 open reading frames, including 5’ untranslated and upstream regions, with single nucleotide polymorphisms highlighted. B. Multiple sequence alignment of sequenced region for KP56 cell line, compared with corresponding regions of Tb927.8.4040 and Tb927.8.4090. Single nucleotide polymorphism indicated with asterisk (*) at position 50, demonstrates integration targeting Tb927.8.4040, not Tb927.8.4090.

## Notes

### Competing Interest Statement

The authors have declared no competing interest.

## REFERENCES

1. Aeschlimann, S., Stettler, P., & Schneider, A. (2023). DNA segregation in mitochondria and beyond: insights from the trypanosomal tripartite attachment complex. Trends in Biochemical Sciences, 48(12), 1058–1070.

2. Albanaz, A. T. S., Carrington, M., Frolov, A. O., Ganyukova, A. I., Gerasimov, E. S., Kostygov, A. Y., … Butenko, A. (2023). Shining the spotlight on the neglected: new high-quality genome assemblies as a gateway to understanding the evolution of Trypanosomatidae. BMC Genomics, 24, 471 (2023).

3. Alfonzo, J. D., Thiemann, O., & Simpson, L. (1997). The mechanism of U insertion/deletion RNA editing in kinetoplastid mitochondria. Nucleic Acids Research, 25(19), 3751–3759.

4. Alsford, S., Turner, D. J., Obado, S. O., Sanchez-Flores, A., Glover, L., Berriman, M., … Horn, D. (2011). High-throughput phenotyping using parallel sequencing of RNA interference targets in the African trypanosome. Genome Research, 21(6), 915–924.

5. Amodeo, S., Bregy, I., & Ochsenreiter, T. (2022). Mitochondrial genome maintenance - the kinetoplast story. FEMS Microbiol Rev. 2023 Nov 1;47(6):fuac047.

6. Armenteros, J. J. A., Salvatore, M., Emanuelsson, O., Winther, O., Von Heijne, G., Elofsson, A., & Nielsen, H. (2019). Detecting sequence signals in targeting peptides using deep learning. Life Sci Alliance. 2019 Sep 30;2(5):e201900429.

7. Benz, C., Dondelinger, F., McKean, P. G., & Urbaniak, M. D. (2017). Cell cycle synchronisation of *Trypanosoma brucei* by centrifugal counter-flow elutriation reveals the timing of nuclear and kinetoplast DNA replication. Sci Rep. 2017 Dec 14;7(1):17599.

8. Billington, K., Halliday, C., Madden, R., Dyer, P., Barker, A. R., Moreira-Leite, F. F., … Gull, K. (2023). Genome-wide subcellular protein map for the flagellate parasite *Trypanosoma brucei*. Nat Microbiol. 2023 Mar;8(3):533-547.

9. Burger, G., & Valach, M. (2018). Perfection of eccentricity: Mitochondrial genomes of diplonemids. IUBMB Life, 70(12), 1197–1206.

10. Cadena, L. R., Gahura, O., Panicucci, B., Zíková, A., & Hashimi, H. (2021). Mitochondrial Contact Site and Cristae Organization System and F1FO-ATP Synthase Crosstalk Is a Fundamental Property of Mitochondrial Cristae. mSphere. 2021 Jun 16;6(3):e0032721.

11. Cadena, L. R., Hammond, M., Tesařová, M., Chmelová, Ľ., Svobodová, M., Durante, I. M., … Lukeš, J. (2024). A novel nabelschnur protein regulates segregation of the kinetoplast DNA in *Trypanosoma brucei*. *BioRxiv*, 2024.03.18.585547.

12. Cymerman, I. A., Chung, I., Beckmann, B. M., Bujnicki, J. M., & Meiss, G. (2008). EXOG, a novel paralog of Endonuclease G in higher eukaryotes. Nucleic Acids Research, 36(4), 1369–1379.

13. Dean, S., Sunter, J., Wheeler, R., Hodkinson, I., Gluenz, E., & Gull, K. (2015). A toolkit enabling efficient, scalable and reproducible gene tagging in trypanosomatids. Open Biology, 5(*1*), 140197– 140197.

14. Desy, S., Schneider, A., & Mani, J. (2012). *Trypanosoma brucei* has a canonical mitochondrial processing peptidase. Molecular and Biochemical Parasitology, 185(2), 161–164.

15. Díaz-Olmos, Y., Batista, M., Ludwig, A., & Marchini, F. K. (2020). Characterising ISWI chromatin remodeler in *Trypanosoma cruzi*. Mem Inst Oswaldo Cruz. 2020;115:e190457.

16. Dibus, N., Korinek, V., & Cermak, L. (2022). FBXO38 Ubiquitin Ligase Controls Centromere Integrity via ZXDA/B Stability. Front Cell Dev Biol. 2022 Jun 23;10:929288. Erratum in: *Front Cell Dev Biol*. 2022 Sep 14;10:1000878.

17. Diez, H., Sarmiento, L., Caldas, M. L., Montilla, M., Thomas, M. D. C., Lopez, M. C., & Puerta, C. (2008). Cellular location of KMP-11 protein in *Trypanosoma rangeli*. *Vector Borne and Zoonotic Diseases (Larchmont*, N.Y.), 8(1), 93–96.

18. Durante, I. M., Butenko, A., Rašková, V., Charyyeva, A., Svobodová, M., Yurchenko, V., … Lukeš, J. (2020). Large-Scale Phylogenetic Analysis of Trypanosomatid Adenylate Cyclases Reveals Associations with Extracellular Lifestyle and Host-Pathogen Interplay. Genome Biology and Evolution, 12(12), 2403–2416.

19. Englund, P. T. (1979). Free minicircles of kinetoplast DNA in *Crithidia fasciculata*. Journal of Biological Chemistry, 254(11), 4895–4900.

20. Fukasawa, Y., Tsuji, J., Fu, S. C., Tomii, K., Horton, P., & Imai, K. (2015). MitoFates: Improved Prediction of Mitochondrial Targeting Sequences and Their Cleavage Sites. Molecular & Cellular Proteomics: MCP, 14(4), 1113.

21. Gannavaram, S., Vedvyas, C., & Debrabant, A. (2008). Conservation of the pro-apoptotic nuclease activity of endonuclease G in unicellular trypanosomatid parasites. Journal of Cell Science, 121(Pt 1), 99–109.

22. Gerasimov, E. S., Gasparyan, A. A., Afonin, D. A., Zimmer, S. L., Kraeva, N., Lukeš, J., … Kolesnikov, A. (2021). Complete minicircle genome of *Leptomonas pyrrhocoris* reveals sources of its non- canonical mitochondrial RNA editing events. Nucleic Acids Research, 49(6), 3354–3370.

23. Hoffmann, A., Käser, S., Jakob, M., Amodeo, S., Peitsch, C., Týč, J., … Ochsenreiter, T. (2018). Molecular model of the mitochondrial genome segregation machinery in *Trypanosoma brucei*. Proceedings of the National Academy of Sciences of the United States of America, 115(8), E1809– E1818.

24. Jensen, R. E., & Englund, P. T. (2012). Network news: the replication of kinetoplast DNA. Annual Review of Microbiology, 66, 473–491.

25. Käser, S., Oeljeklaus, S., Týc, J., Vaughan, S., Warscheid, B., & Schneider, A. (2016). Outer membrane protein functions as integrator of protein import and DNA inheritance in mitochondria. Proceedings of the National Academy of Sciences of the United States of America, 113(31), E4467–E4475.

26. Kostygov, Alexei Y., Karnkowska, A., Votýpka, J., Tashyreva, D., Maciszewski, K., Yurchenko, V., & Lukeš, J. (2021). Euglenozoa: taxonomy, diversity and ecology, symbioses and viruses. Open Biol. 2021 Mar;11(3):200407.

27. Kostygov, Alexei Yu, Albanaz, A. T. S., Butenko, A., Gerasimov, E. S., Lukeš, J., & Yurchenko, V. (2024). Phylogenetic framework to explore trait evolution in Trypanosomatidae. Trends in Parasitology, 40(2), 96–99.

28. Li, S. J., Zhang, X., Lukeš, J., Li, B. Q., Wang, J. F., Qu, L. H., … Lun, Z. R. (2020). Novel organization of mitochondrial minicircles and guide RNAs in the zoonotic pathogen *Trypanosoma lewisi*. Nucleic Acids Research, 48(17), 9747–9761.

29. Li, Z., & Wang, C. C. (2008). KMP-11, a basal body and flagellar protein, is required for cell division in *Trypanosoma brucei*. Eukaryotic Cell, 7(11), 1941–1950.

30. Lindsay, M. E., Gluenz, E., Gull, K., & Englund, P. T. (2008). A new function of *Trypanosoma brucei* mitochondrial topoisomerase II is to maintain kinetoplast DNA network topology. Molecular Microbiology, 70(6), 1465–1476.

31. Liu, B., Wang, J., Yaffe, N., Lindsay, M. E., Zhao, Z., Zick, A., … Englund, P. T. (2009). Trypanosomes Have Six Mitochondrial DNA Helicases with One Controlling Kinetoplast Maxicircle Replication. Molecular Cell, 35(4), 490–501.

32. Lukes, J., Guilbride, D. L., Voty, J., Zíkova, A., Benne, R., & Englund, P. T. (2002). Kinetoplast DNA Network : Evolution of an Improbable Structure. Eukaryot Cell. 2002 Aug;1(4):495-502.

33. Lukeš, J., Speijer, D., Zíková, A., Alfonzo, J. D., Hashimi, H., & Field, M. C. (2023). Trypanosomes as a magnifying glass for cell and molecular biology. Trends in Parasitology, 39(11), 902–912.

34. Marques, C. A., Ridgway, M., Tinti, M., Cassidy, A., & Horn, D. (2022). Genome-scale RNA interference profiling of *Trypanosoma brucei* cell cycle progression defects. Nat Commun. 2022 Sep 10;13(1):5326.

35. Maslov, D. A. (2019). Separating the wheat from the chaff: RNA editing and selection of translatable mRNA in trypanosome mitochondria. Pathogens. 2019 Jul 18;8(3):105.

36. Meehan, J., McDermott, S. M., Ivens, A., Goodall, Z., Chen, Z., Yu, Z., … Cruz-Reyes, J. (2023). Trypanosome RNA helicase KREH2 differentially controls non-canonical editing and putative repressive structure via a novel proposed “bifunctional” gRNA in mRNA A6. Nucleic Acids Research, 51(13), 6944–6965.

37. Moloney, N. M., Barylyuk, K., Tromer, E., Crook, O. M., Breckels, L. M., Lilley, K. S., … MacGregor, P. (2023). Mapping diversity in African trypanosomes using high resolution spatial proteomics. Nat Commun. 2023 Jul 21;14(1):4401.

38. Panigrahi, A. K., Ogata, Y., Zíková, A., Anupama, A., Dalley, R. A., Acestor, N., … Stuart, K. D. (2009). A comprehensive analysis of *Trypanosoma brucei* mitochondrial proteome. Proteomics. 2009 Jan;9(2):434-50.

39. Peikert, C. D., Mani, J., Morgenstern, M., Käser, S., Knapp, B., Wenger, C., … Warscheid, B. (2017). Charting organellar importomes by quantitative mass spectrometry. Nat Commun. 2017 May 9;8:15272.

40. Poon, S. K., Peacock, L., Gibson, W., Gull, K., & Kelly, S. (2012). A modular and optimized single marker system for generating *Trypanosoma brucei* cell lines expressing T7 RNA polymerase and the tetracycline repressor. Open Biol. 2012 Feb;2(2):110037.

41. Povelones, M. L. (2014). Beyond replication: division and segregation of mitochondrial DNA in kinetoplastids. Mol Biochem Parasitol. 2014 Aug;196(1):53-60.

42. Pusnik, M., Mani, J., Schmidt, O., Niemann, M., Oeljeklaus, S., Schnarwiler, F., … Schneider, A. (2012). An essential novel component of the noncanonical mitochondrial outer membrane protein import system of trypanosomatids. Molecular Biology of the Cell, 23(17), 3420–3428.

43. Pyrih, J., Hammond, M., Alves, A., Dean, S., Sunter, J. D., Wheeler, R. J., … Lukeš, J. (2023). Comprehensive sub-mitochondrial protein map of the parasitic protist *Trypanosoma brucei* defines critical features of organellar biology. Cell Rep. 2023 Sep 26;42(9):113083.

44. Redmond, S., Vadivelu, J., & Field, M. C. (2003). RNAit: An automated web-based tool for the selection of RNAi targets in *Trypanosoma brucei*. Mol Biochem Parasitol. 2003 Apr 25;128(1):115-8.

45. Rosamond, J. (1981). Purification and properties of an endonuclease from the mitochondrion of *Saccharomyces cerevisiae*. Eur J Biochem. 1981 Dec;120(3):541-6.

46. Schindelin, J., Arganda-Carreras, I., Frise, E., Kaynig, V., Longair, M., Pietzsch, T., … Cardona, A. (2012, July). Fiji: An open-source platform for biological-image analysis. Nat Methods. 2012 Jun 28;9(7):676-82.

47. Schneider, A., & Ochsenreiter, T. (2018). Failure is not an option - mitochondrial genome segregation in trypanosomes. J Cell Sci. 2018 Sep 17;131(18):jcs221820.

48. Sherwin, T., & Gull, K. (1989). The cell division cycle of *Trypanosoma brucei brucei*: timing of event markers and cytoskeletal modulations. Philos Trans R Soc Lond B Biol Sci. 1989 Jun 12;323(1218):573-88.

49. Shlomai, J. (2004). The structure and replication of kinetoplast DNA. Curr Mol Med. 2004 Sep;4(6):623- 47.

50. Stuart, K., Brun, R., Croft, S., Fairlamb, A., Gürtler, R., McKerrow, J., … Tarleton, R. (2008). Kinetoplastids: related protozoan pathogens, different diseases. J Clin Invest. 2008 Apr;118(4):1301- 10.

51. Stuart, K. D., Schnaufer, A., Ernst, N. L., & Panigrahi, A. K. (2005). Complex management: RNA editing in trypanosomes. Trends Biochem Sci. 2005 Feb;30(2):97-105.

52. Votýpka, J., Kostygov, A. Y., Kraeva, N., Grybchuk-Ieremenko, A., Tesařová, M., Grybchuk, D., … Yurchenko, V. (2014). Kentomonas gen. n., a new genus of endosymbiont-containing trypanosomatids of Strigomonadinae subfam. n. Protist, 165(6), 825–838.

53. Wang, J., Englund, P. T., & Jensen, R. E. (2012). TbPIF8, a *Trypanosoma brucei* protein related to the yeast Pif1 helicase, is essential for cell viability and mitochondrial genome maintenance. Molecular Microbiology, 83(3), 471–485.

54. Wang, Z., & Englund, P. T. (2001). RNA interference of a trypanosome topoisomerase II causes progressive loss of mitochondrial DNA. EMBO J. 2001 Sep 3;20(17):4674-83.

55. Wenger, C., Oeljeklaus, S., Warscheid, B., Schneider, A., & Harsman, A. (2017). A trypanosomal orthologue of an intermembrane space chaperone has a non-canonical function in biogenesis of the single mitochondrial inner membrane protein translocase. PLoS Pathog. 2017 Aug 21;13(8):e1006550.

